# Changes in interictal pretreatment and posttreatment EEG in childhood absence epilepsy

**DOI:** 10.1101/699868

**Authors:** Pawel Glaba, Miroslaw Latka, Małgorzata Krause, Marta Kuryło, Wojciech Jernajczyk, Wojciech Walas, Bruce J. West

## Abstract

Spike and wave discharges (SWDs) are the characteristic manifestation of childhood absence epilepsy (CAE). It has long been believed that they unpredictably emerge from otherwise almost normal interictal EEG. Herein, we demonstrate that pretreatment closed-eyes theta and beta EEG wavelet powers of CAE patients (20 girls and 10 boys, mean age 7.4 *±* 1.9 years) are much higher than those of age-matched controls at multiple sites of 10-20 system. For example, at C4 site, we observed a 91% and 62% increase in power of theta and beta rhythms, respectively. We were able to compare the baseline and posttreatment wavelet power in 16 patients. The pharmacotherapy brought about a statistically significant decrease in delta and theta wavelet power in all the channels, e.g. for C4 the reduction was equal to 45% (delta) and 65% (theta). We also observed a less pronounced attenuation of posttreatment beta rhythm in several channels. We hypothesize that the increased theta and beta powers result from cortical hyperexcitability and propensity for epileptic spikes generation, respectively. We argue that the distinct features of CAE wavelet power spectrum may be used to define an EEG biomarker which could be used for diagnosis and monitoring of patients.

## Introduction

Childhood absence epilepsy (CAE) is the most common pediatric epileptic syndrome. It has a prevalence of 10–15% in childhood epilepsies and an incidence of 1.3 to 6 per 100,000 in children under the age of 16 years (Covanis, 2010). The disorder is most likely multifactorial, resulting from interactions between genetic and acquired factors (Gallentine and Mikati, 2012).

The ictal EEG of a typical absence seizure demonstrates rhythmic ~3 Hz bilateral, synchronous and symmetrical spike and wave discharges (SWDs) with median duration of approximately 10 seconds, which may appear several times per day, sometimes as often as dozens times per day (Schomer and Lopes da Silva, 2018). It has long been believed that SWDs are unpredictable and emerge from otherwise almost normal interictal EEG (Lopes da Silva *et al.*, 2003). Interictal EEG abnormalities include sparse fragments of SWDs and focal discharges as well as posterior bilateral delta activity (Sadleir *et al.*, 2006).

The response to monotherapy (valproic acid, ethosuximide, lamotrigine) is generally good. As high as 75-85% of treated patients are seizure free and have a normal EEG (Covanis, 2010). The rest usually respond to the combination of drugs such as valproic acid and ethosuximide. The pharmacological seizure control with acceptable side effects is achieved for slightly more than half of the children. The relationship between pretreatment EEG and response to therapy is unknown (Dlugos *et al.*, 2013). In general CAE carries a good prognosis. Seizures spontaneously cease with ongoing maturation.

The cortical focus theory postulates that an absence seizure originates from a focus in the somatosensory cortex which is in a specific state conducive to seizure propagation. At the beginning of the seizure, it is the cortex that drives the thalamus, but prominent generalized spike-wave discharges result from their subsequent mutual interaction (Meeren *et al.*, 2005*a*). Taking into account the principal role of the cortex in absence seizure generation, we hypothesize that in CAE, interictal EEG manifests distinct features that can be detected even in routine, rest EEG recording.

## Materials and methods

We retrospectively analyzed a routine pretreatment video EEG recording of n=30 CAE patients (20 girls and 10 boys) with mean age of 7.4 *±* 1.9 years. The analysis was approved by the board of the T. Marciniak Hospital in Wroclaw, Poland. The CAE epilepsy syndrome was established on the basis of history, age at onset, clinical and EEG findings as well as neuroimaging. In this cohort, we observed 173 seizures, 6 *±* 3 per patient on average. The median duration of seizure was 12 *±* 4 seconds. In 9 cases, the first seizure occurred during hyperventilation, on average 130 *±* 55 seconds after its onset. The control group (15 girls and 15 boys) was age matched (7.3 *±* 1.9 years) to the controls. EEG was recorded using Elmiko Digitrack system with a BRAINTRONICS B.V. ISO-1032CE amplifier. The sampling frequency was equal to 250 Hz. The 10-20 international standard was used to position 19 Ag/AgCl electrodes (impedances were below 5 *k*Ω). The ground electrode was placed at the patients’ forehead. The average reference electrode montage was used for time-frequency calculations. All the EEG recordings were performed between 2008 and 2018 by the same certified EEG technician.

After diagnosis, the pharmacotherapy of 16 out of 30 patients was administered by the outpatient neurology department of T. Marciniak Hospital. Valproic acid (two daily dosages of 244 *±* 58 mg and 311 *±* 111 mg) was used in 73% of the cases and 18% of the patients were treated with ethosuximide (two daily dosages of 250 mg). The combination of valproic acid (250 mg twice a day) and ethosuximide (150 mg twice a day) was used in 9% of the patients. The treatment response was evaluated with routine EEG with hyperventilation and photostimulation.

For each patient, we extracted a closed-eyes (CE) motion artifact free pretreatment EEG segment that preceded the first seizure. The average length of these segments was: 84 *±* 32 s. For the controls, the average length of selected EEG segments was equal to 55 *±* 27 s.

We also analyzed the most recent follow-up EEG which on average was recorded 1.5 *±* 2 years after the video EEG used for diagnosis. The average length of CE segments selected for analysis from the follow-up EEG was 28 *±* 15 s. There were no seizures in the follow-up EEGs.

We calculated the continuous wavelet transform (CWT) of EEG using the complex Morlet mother function with the center frequency *f*_*c*_=1 and bandwidth *f*_*b*_=1.8 (Latka *et al.*, 2003). For these parameters, the transform was computed for three frequencies *f:* 6, 10, and 18 Hz. The first two frequencies roughly correspond to the centers of theta and alpha bands. Fig. 1 elucidates the rationale for choosing 18 Hz for beta rhythm analysis. The parameters of the wavelet calculations were tuned for detection of epileptic spikes. In other words, we focus on this part of the beta band which may be involved in the genesis of epileptic spikes. The spectacular increase of beta wavelet power in the vicinity of the epileptic spikes in Fig. 1 can be employed in the simple but highly effective SWD detector which we describe elsewhere. The wavelet with *f*_*c*_=1 and bandwidth *f*_*b*_=1.5 calculated for frequency 3 Hz was used to quantify delta waves. We analyzed the statistical properties of instantaneous wavelet power *w*(*f, t*_0_) (square of the modulus of complex wavelet coefficients) and temporal average *W* (*f*) =*< w*(*f, t*_0_) *>*_*t0*_ over the selected EEG segment.

**Figure 1.**
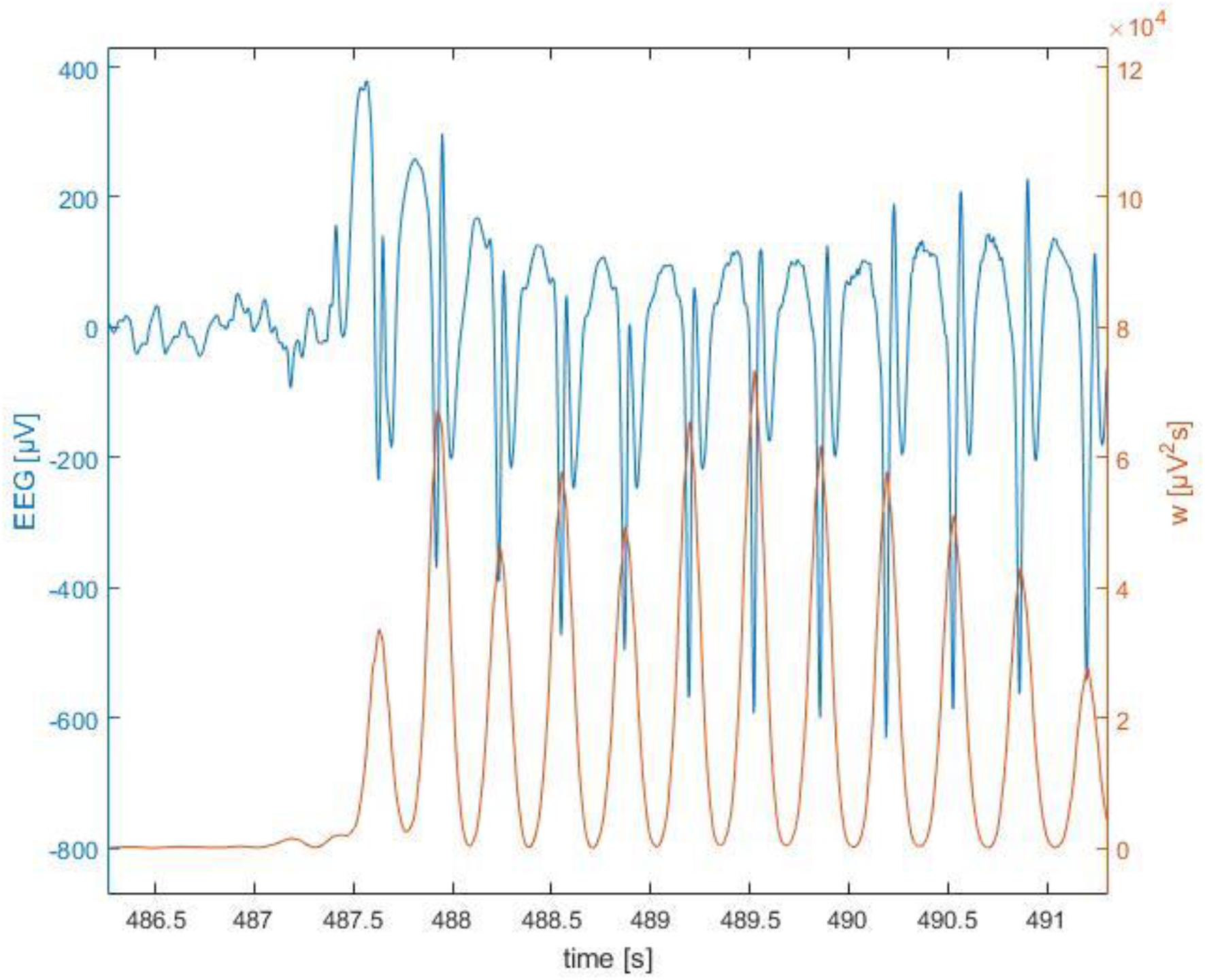
Example of childhood absence epilepsy seizure. The presented section of EEG (blue curve) was extracted from channel Fp1 of a seven-year-old girl (patient BZ). The instantaneous power of the complex Morlet wavelet (*f*_*c*_=1,*f*_*b*_=1.8) calculated for 18 Hz was drawn with the red curve.

We used the Shapiro-Wilk test to determine whether the analyzed data were normally distributed. The significance threshold for all the statistical tests was set to 0.05. We compared the pretreatment and posttreatment values of *W* (*f*) as well as the values of controls with the Kruskal-Wallis test (with Tukey’s post hoc comparisons). The area under the receiver operating characteristic curve (AUC) was used to quantify differences in *W* (*f)* between the patients and controls. AUC was also calculated for pretreatment and posttreatment values of *W*. The values of AUC, sensitivity and specificity were reported for selected channels.

### Data availability

The authors confirm that the data supporting the findings of this study are available within the article and its supplementary material.

## Results

Fig. 2 shows that the pretreatment closed-eyes theta and beta EEG wavelet powers of CAE patients are much higher than those of age-matched controls at multiple sites of 10-20 system. For theta band, we observed a 91% increase in patients’ wavelet power *W* at C4 site (345 *±* 286 *µV* ^2^*s* vs 181 *±* 120 *µV* ^2^*s*, *p* = 8 *×*10^*−*3^, AUC=0.73, sensitivity=0.67, specificity =0.67). For beta band, the power in the patients was 62% greater than in the controls (23 *±* 10 *µV* ^2^*s* vs 14*±* 8 *µV* ^2^*s*, *p* = 7 *×* 10^*−*3^, AUC=0.80, sensitivity=0.73, specificity=0.74). There was no statistically significant difference between the patients and the controls for alpha rhythm. For delta band, the increased wavelet power in the patients was observed only at the occipital channels.

**Figure 2.**
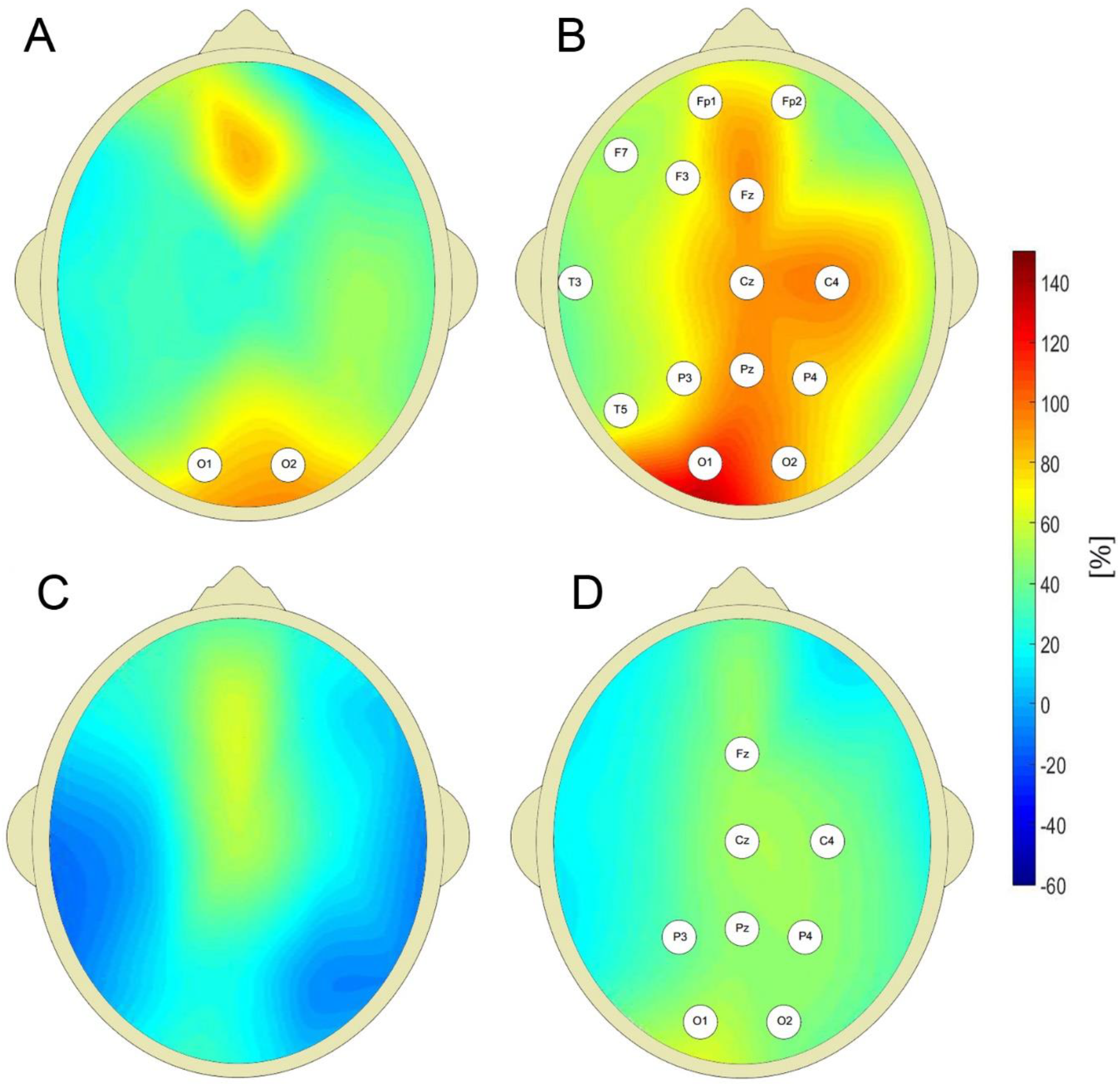
Percentage differences between wavelet power of closed eyes interictal EEG of CAE patients and the controls (with respect to controls) for: delta (A), theta (B), alpha (C), and beta (D) rhythms. The EEG channel labels indicate sites for which the differences were statistically significant.

In Fig. 3 we present the probability density function (PDF) of instantaneous theta and beta wavelet power *w(f,t*_*0*_) in channel C4 for the controls (light green) and the patients (dark green). For both cohorts, we aggregated the values of *w(f,t*_*0*_) from all their members. It is apparent that the tail of the PDF is much longer for the pretreatment cohort. The insets in both subplots show the boxplots of time-averaged values of the pretreatment and posttreatment wavelet power *W* as well as that of the controls. It is apparent from Fig. 3A that the pharmacotherapy suppressed the elevated pretreatment theta power that dropped 65% from 345 *±* 286 *µV* ^2^*s* to 120*±* 106 *µV*^2^*s* with posthoc *p*=5*×* 10^*−*5^ (AUC=0.87, sensitivity=0.90, specificity =0.75). The patients’ posttreatment C4 theta power was not different from that of the controls (*p*=0.16). Fig. 3B shows that the influence of treatment on C4 beta power was weaker. In that case the power was reduced 37% from 23 *±* 10 *µV*^2^*s* to 14 *±* 12 *µV*^2^*s* with post hoc *p*=7*×* 10^*−*3^ (AUC=0.74, sensitivity=0.80, specificity =0.63). While the posttreatment median of *W* was not different from that of the controls, the width of the patients’ distribution was clearly greater.

**Figure 3.**
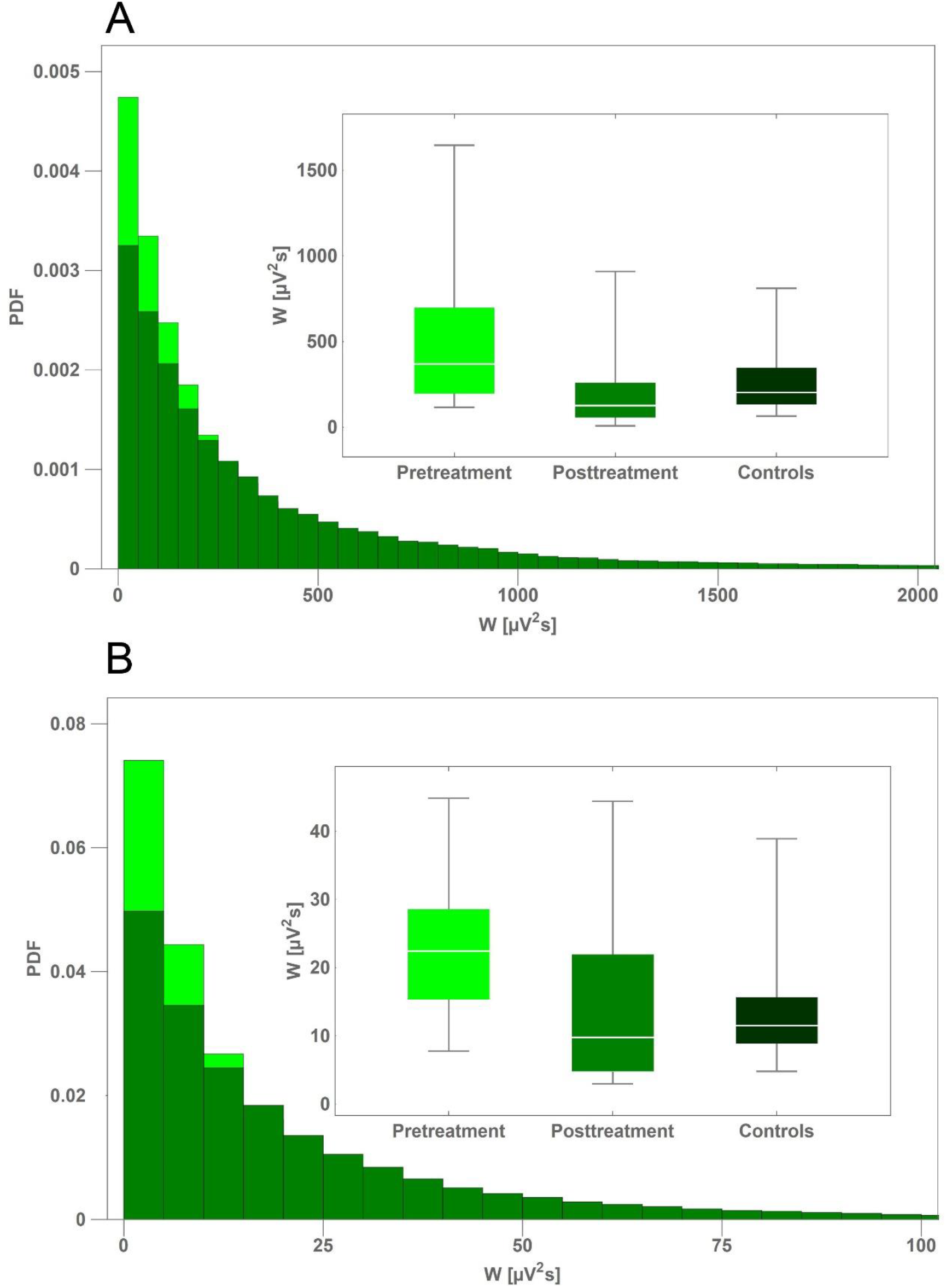
Comparison of probability density function (PDF) of instantaneous wavelet power of pretreatment EEG of CAE patients (light green) and the controls (dark green) for: theta (A) and beta (B) rhythms. PDFs were calculated from the closed-eyes segments extracted from C4 channel. The insets in both subplots show the boxplots of time-averaged values of the pretreatment and posttreatment wavelet power of as well as that of the controls.

One can see in Fig. 4 that pharmacotherapy primarily attenuates the low-frequency part of EEG spectrum. For all the channels, the values of posttreatment delta and theta power are smaller than the pretreatment ones. In particular, for delta band, *W* for C4 channel fell 45% from 394 *±* 280 *µV* ^2^*s* to 218 *±* 328 *µV* ^2^*s* (*p* = 7 *×* 10^*−*3^, AUC=0.90, sensitivity=0.75, specificity=0.84). In sharp contrast, there were only 2 and 7 channels with the reduced power for alpha and beta band, respectively. The fact that there is no increase of posttreatment alpha wavelet power implies that the attenuation of theta rhythm does not result from EEG maturation but is caused by treatment.

**Figure 4.**
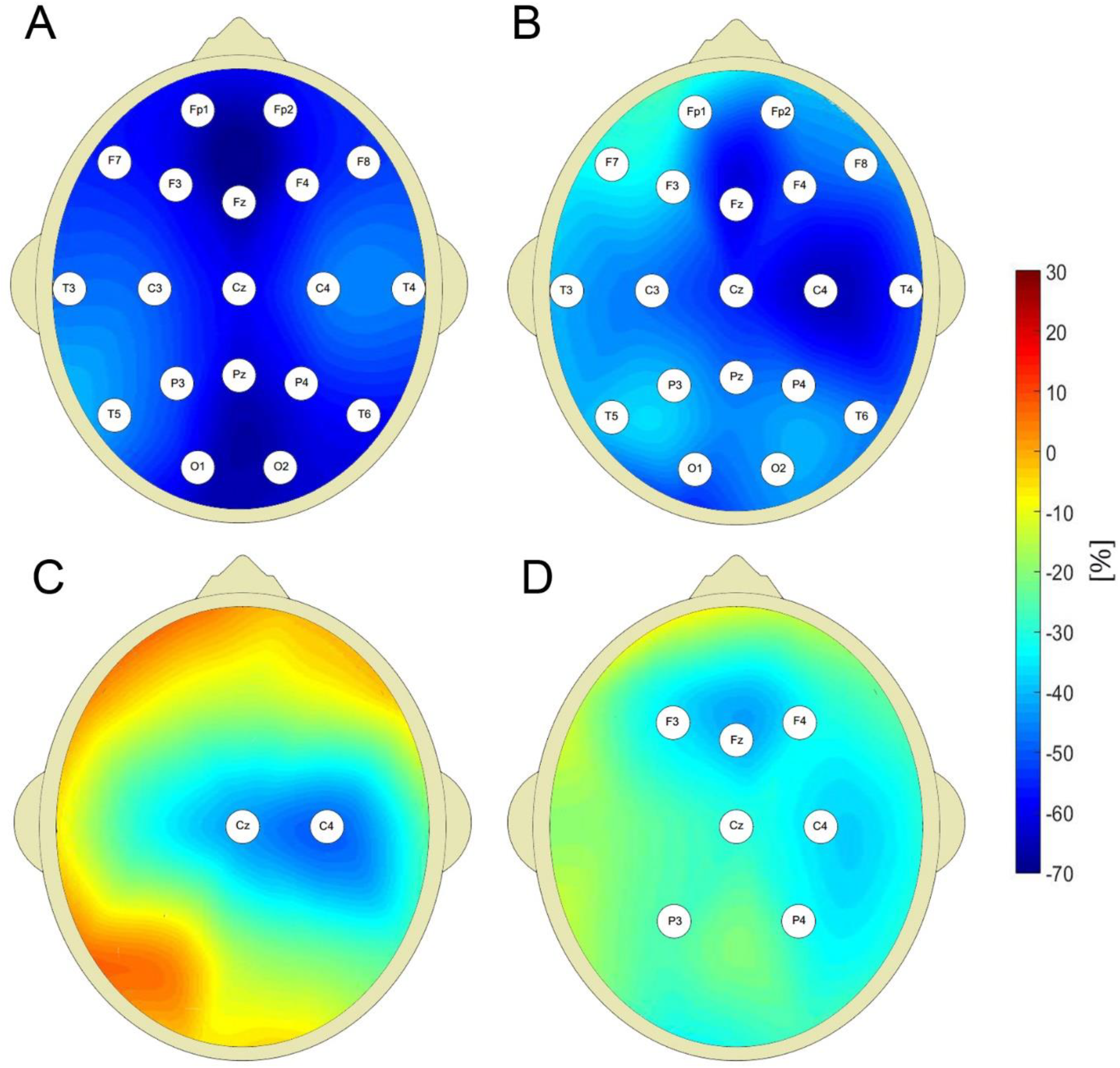
Percentage change in posttreatment closed eyes wavelet power of CAE patients with respect to the baseline for: delta (A), theta (B), alpha (C), and beta (D) rhythms. The EEG channel labels indicate sites for which the differences were statistically significant.

## Discussion

GAERS and the WAG/Rij rodent models have paved the way for the understanding of pathophysiology of human CAE seizures (Akman *et al.*, 2010). Lüttjohann and Van Luijtelaar reviewed the animal studies and translation of research results from rodent models to humans (Lüttjohann and Van Luijtelaar, 2015). According to the cortical focus theory of (Meeren *et al.*, 2005*b*) the genesis of absence seizure starts with a single spike which rapidly spreads over the hyperexcited cortex. The formation of prominent SWDs is possible only due to the interaction with thalamus which acts as a resonant circuitry. The rapid generalization of the spike-wave activity over the cortex is due to short-range intracortical fibers and to a subpopulation of cells that have long-range association fibers. These fibers run under the cortex in the white matter, making extensive connections with other cortical areas.

In rats, SWDs emerge from the cortical focus which is located either within the perioral somatosensory cortex (WAG/Rij) or the secondary somatosensory cortex (GAERS) and subsequently propagate via the corticothalamocortical loop. Polack *et al.* demonstrated that blockade of action potential discharge and synaptic activities in facial somatosensory cortical neurons (by topical application of tetrodotoxin) prevents the formation of SWDs (Polack *et al.*, 2009). In contrast, pharmacological inhibition of a remote motor cortical region does not suppress ictal activities. Westmijse *et al.* discovered, using a beamforming source localization technique, that in humans the sources of spikes from a train of SWDs were at the frontal lateral, central and medial parietal cortices (Westmijse *et al.*, 2009). The involvement of the thalamus in the generation of SWDs was demonstrated in combined EEG-fMRI (Granert *et al.*, 2008; Moeller *et al.*, 2010) and MEG (Tenney *et al.*, 2013) studies. The differences between rat and human data are in the frequencies of SWDs (6-11 Hz vs 2.5-4 Hz) and the location of the early local cortical activity which is quite variable in humans. The location may even change during a seizure (Westmijse *et al.*, 2009) and is predominantly, but not exclusively, located in the frontal-central/parietal areas. The low variability of the position of cortical focus in the rats can easily be explained by the fact that both epileptic strains are fully inbred, and the animals are homozygous.

Herein, we found that pretreatment closed-eyes theta wavelet power of CAE patients was much higher than that of age-matched controls at multiple sites of 10-20 system. This result should not come as a surprise. In the late 1960s Doose *et al.* argued that strong rhythmic theta activity was an age-dependent electroencephalographic expression of a genetic disposition to convulsions, see a review paper (Doose and Baier, 1988) for a complete list of references. This hypothesis was noted, along with the opposing view, by the authors of the classic text on encephalography (Schomer and Lopes da Silva, 2018). To the best of our knowledge, initial observations have not been followed up on using quantitative EEG analysis (Doose *et al.*, 2003). The more recent research provided new evidence for the role of theta rhythm in epilepsy. Douw *et al*. in their MEG studies found that theta band brain connectivity and network topology is altered in epilepsy which developed in brain tumor patients (Douw *et al.*, 2010). Milikovsky *et al*. discussed the properties of theta dynamics in the animal model of postinjury epilepsy (Milikovsky *et al.*, 2017). Sitnikova and van Luijtelaar found that the increased delta and theta power preceded SWDs in WAG/Rij rats (Sitnikova and van Luijtelaar, 2009).

We interpret the results presented herein as evidence in support of the following postulates. We hypothesize that the increased theta rhythm power in CAE patients is a manifestation of cortical hyperexcitability (Vestal and Blumenfeld, 2010). Since strong theta rhythm may be found in up to 15% of healthy children (Doose and Baier, 1988), cortical hyperexcitability may be a necessary but not a sufficient prerequisite for CAE. The increased beta power may reflect propensity for epileptic spikes generation – spikes that triggers absence seizures. This interpretation is corroborated by the recent work of Sorokin *et al*. who found that in rats absence seizure susceptibility correlates with pre-ictal beta oscillations (Sorokin *et al.*, 2018). It is worth pointing out that as we detune the parameters of the analyzing wavelet away from the values optimal for spike detection, the differences in beta power between the CAE patients and controls disappear.

As with all epileptic syndromes, once the diagnosis of CAE is confirmed, only the absence or recurrence of seizures provides an indication as to whether anti-epileptic drugs (AEDs) have had any effect. We argue that the distinct features of CAE wavelet power spectrum may be used to define an EEG biomarker which could be used for diagnosis and the long-term monitoring of patients. The most apparent applications of such monitoring would be in the assessment of the efficacy of pharmacotherapy and its duration.

Over the last three decades, transcranial magnetic stimulation (TMS) has become a principal tool for accessing cortical excitability associated with epilepsy (Bauer *et al.*, 2014). However, there are essentially no TMS studies of drug-naive patients with CAE (Kessler, 2016). The barriers to the inclusion of children from such studies stem not only from the very nature of this experimental technique, e.g. its duration or immobilization of head during measurement, but also from a few fundamental reasons. For example, it is not clear whether the motor cortex physiology is a reliable marker of seizure susceptibility in children with less mature brain networks. Moreover, abnormalities in TMS markers of cortical excitability are not specific to epilepsy and may observed, among others, in ADHD, a common comorbid condition in children with epilepsy (Reilly *et al.*, 2017). Future research should verify whether measurement of theta band power in CAE patients could be used in lieu of TMS. In particular, the previous TMS studies (Wright *et al.*, 2006; Badawy *et al.*, 2009) revealed that increased excitability preceded epileptic seizures. Thus, the question arises as to whether temporal changes in cortical excitability occur in CAE.

The pharmacological seizure control with acceptable side effects is achieved for slightly more than half of the children with CAE. Herein we found that pharmacotherapy most effectively suppresses low-frequency EEG oscillations. Quantitative EEG analysis offers a unique opportunity to design a treatment which would more selectively attenuate the pathological theta and/or beta rhythms. The availability of low-cost, Internet connected personal EEG devices paves the way for home monitoring of patients which would facilitate drug titration and termination of pharmacotherapy.

